# Comparing electromyography, accelerometry, and visual inspection to assess the resting motor threshold for transcranial magnetic stimulation

**DOI:** 10.1101/2025.06.04.657787

**Authors:** Gautier Hamoline, Antoine E Lunardi, Marcos Moreno-Verdú, Baptiste M Waltzing, Elise E Van Caenegem, Siobhan MacAteer, Robert M Hardwick

## Abstract

**Introduction:** Electromyography (EMG) remains the gold standard for estimating the Resting Motor Threshold (RMT) in Transcranial Magnetic Stimulation (TMS) studies, but its cost and limited accessibility often lead researchers to use visual inspection (VIS). However, VIS may introduce variability and systematic bias. Accelerometry (ACC) offers a cost-effective, objective alternative to capture TMS-evoked responses.

**Objective:** To compare the RMT as estimated using EMG, ACC, and VIS.

**Methods:** Five participants underwent TMS while EMG, ACC, and video recordings were collected. Separately, 64 observers judged hand movement in videos to estimate RMT via VIS. RMTs were compared across the three methods using Bayesian model comparison, Bland-Altman analyses, and Intraclass Correlation Coefficients (ICCs).

**Results:** RMTs estimated via EMG were lower than those obtained using either ACC or VIS. Compared to EMG, VIS tended to overestimate RMT (mean bias = 5.23%, 95%CI = [1.00–11.00]), while ACC and VIS estimates were more closely aligned (mean bias = 0.43%, 95%CI = [–4.00 – 5.00]). ICC (2,1) values indicated moderate reliability for VIS vs EMG (mean = 0.580, 95%CI = [0.389 – 0.748]), and good-to-excellent reliability for VIS vs ACC (mean = 0.845). However, bootstrapped 95% confidence intervals identified significant variability in the estimates provided by visual inspection, ranging from +1 to +11 for VIS vs EMG, but as low as −4 to +5 for VIS vs ACC.

**Conclusions:** EMG remains the most sensitive technique for estimating the RMT, but when EMG is not feasible, accelerometery provides a quantifiable, more objective, and less variable alternative than visual inspection.

**Highlights:**

- Visual inspection introduces observer-related variability in RMT estimation
- Visual inspection & accelerometry systematically overestimate RMT compared to EMG
- EMG remains the most sensitive method for estimating RMT
- Accelerometry is an objective alternative to visual inspection if EMG is unavailable

## 1. Introduction

Research on the effects of Transcranial Magnetic Stimulation (TMS) has largely focused on the primary motor cortex of the human brain (1). When an area of the primary motor cortex linked to a target muscle is stimulated, it can activate the corticospinal pathway leading to muscle contractions, and potentially induce movements in that muscle. By employing electromyography (EMG), the resulting muscle activity from this stimulation can be recorded as a ‘Motor Evoked Potential’ or MEP (2). Among the features of the MEP, the most commonly analysed is its amplitude, often measured from peak to peak in microvolts (µV). Measuring MEP amplitudes is central to defining the intensity of TMS that is delivered in most TMS studies, which is typically defined based on the Resting Motor Threshold (RMT) — the minimal stimulator output necessary to produce MEPs with an amplitude of ≥50µV in at least 5 out of 10 trials.

Although much of the early research on TMS has focused on fundamental motor responses, over time, TMS has garnered significant attention for its wide-ranging applications across diverse fields. In neurology, TMS has been used to examine conditions such as multiple sclerosis (3–7), stroke (8,9), epilepsy (10,11), Parkinson’s disease (12–15), dystonia (16–18) and myopathy (19–21). The application of repetitive TMS has demonstrated substantial promise in treating psychiatric disorders, leveraging its ability to modulate neural activity in targeted brain regions. Conditions such as major depressive disorder (MDD) (22–24), obsessive-compulsive disorder (OCD) (23,25–27), anxiety(28), post-traumatic stress disorder (PTSD)(29,30) and schizophrenia (31,32) have all benefited from TMS interventions.

Additionally, its efficacy extends to behavioral disorders, including addiction (33), attention-deficit/hyperactivity disorder (ADHD) (34), and eating disorders (35). Beyond these possible therapeutic applications, TMS has also proven to be a valuable tool in cognitive research, to improve our understanding of cognitive processes such as mental imagery, decision making and action preparation (36). Thus, while TMS was initially developed to probe motor system function through the measurement of MEPs, it has since evolved into a versatile tool with broad applications.

Regardless of the exact field of study, EMG remains the gold standard for assessing TMS responses. However, EMG presents several drawbacks. The initial cost of EMG equipment such as apparatus and amplifiers, combined with recurring expenses for consumables like electrodes, electrode gel, and cleaning alcohol, makes it a relatively expensive technique (37). Additionally, precise electrode placement requires understanding of muscle anatomy and physiology (38). Preparatory steps, including skin cleaning, hair removal, and verifying the clarity of the signal, all add to the time required to create optimal recordings using this method (39). Despite its widespread use among research teams investigating motor control and neurophysiology, many laboratories employ EMG solely to evaluate the RMT (40–42). However, due to the costs and preparation time associated with EMG, some groups instead measure the “visually determined” RMT. This approach defines RMT as the lowest stimulator output that generates a visible movement (usually of the hand) in at least 5 out of 10 trials (43). While more accessible and less expensive than using EMG, the visual inspection method is inherently subjective (43) and therefore subject to inter- and intra-individual variability; moreover, it focuses on identifying the presence/absence of movement without providing other quantifiable measures (e.g. the speed or magnitude of the induced movement). Furthermore, previous reports provide conflicting accounts of the relationship between measures collected using visual inspection and EMG, with research indicating the visually defined threshold can differ from that identified using EMG by between –10 to +2% of maximal stimulator output (43,44). This is notable as under- or over-estimating the RMT could in turn lead to the application of stimulation intensities that are too low to be effective, or that may exceed TMS safety guidelines (45). Together, these limitations underscore the need to investigate alternative methods to EMG that are both cost-effective and efficient, but also allow precise quantification of TMS evoked responses.

Accelerometry emerges as a promising solution to address the issues with EMG and visual inspection in assessing TMS responses. Specifically, accelerometry is more cost-effective than EMG, requiring smaller initial outlays and incurring lower recurring expenses. Accelerometry also enables researchers to objectively quantify TMS responses, and is therefore likely to be more reliable than the subjective assessments provided by visual inspection. Past studies have used accelerometry to examine how TMS-evoked movement direction and magnitude change with training (46–48). Recently, our laboratory has explored using accelerometry as a tool to measure different characteristics of TMS evoked responses; although accelerometry tends to overestimate the RMT, comparisons of measurements of MEP amplitudes and accelerometer jerk readings were highly correlated, demonstrating this measurement tool shows promising potential (49).

In this paper, the aim is to investigate and compare the measurement of the Resting Motor Threshold (RMT) using three different methods: electromyography (EMG), accelerometry (ACC), and visual inspection (VIS). By evaluating the accuracy, reliability, and practical application of each technique, the present study will determine their respective strengths and limitations in capturing the RMT.

## 2. Materials and Methods

All code and data are available on the following OSF repository : https://osf.io/2mpxf/.

### 2.1. General Design

The study leveraged a mixed-model design to examine data from a total sample of 105 participants. The final sample after application of exclusion criteria (see below) included 69 participants. This final sample comprised two groups of participants.

Data from Group 1 provided direct recordings of responses to TMS. Data collection involved recording data from five participants using three methods: electromyography, accelerometry, and video recordings of the hand during TMS (i.e. recordings demonstrating whether the intensity of TMS was sufficient to produce movement or not).

Data from Group 2 then focused on the visual determination of the resting motor threshold by individual observers. Participants in this second group observed video recordings taken from the first group, providing responses to indicate whether they observed a movement in each trial in response to TMS.

### 2.2. Group 1: Recording of responses to TMS (EMG, Accelerometry, and Video)

#### 2.2.1. Participants

Five participants (2 females, 3 males), all right-handed, with an average age of 24.2 years (ranging from 21 to 29 years, SD = ± 2.31) took part in this part of the study. The safety screening checklist recommended by Lefaucheur et al. (50) was administered prior to stimulation to confirm participant eligibility and ensure safe application for the use of TMS. The study was approved by the Ethics Committee Saint-Luc Hospital, UCLouvain (2020/30JUL/389 - DNMT-TMS – amendment 2), and each participant provided written informed consent prior to the experiment. Participants also consented to video recordings of their hands to be presented to other participants (see below).

#### 2.2.2. Transcranial Magnetic Stimulation (TMS)

TMS was administered using a monophasic Magstim 200^2^ stimulator with a figure-of-eight coil (2 x Ø70 mm) directed posteriorly and toward the ipsilateral side with a random interval of 4 to 6 seconds between each stimulation. To ensure precise targeting, participants wore a head tracker (NT-103) and the Visor2 neuronavigation system (ANT Neuro, Netherlands) was used with the standard brain model, which provided real-time guidance for accurate coil positioning.

#### 2.2.3. Electromyography (EMG)

EMG was utilized to record Motor Evoked Potentials (MEPs) from the First Dorsal Interosseus (FDI) muscle of the index finger on the dominant hand. Before placing the electrodes, the skin was cleaned with an alcohol solution and shaved if necessary to remove hair and enhance conductivity. Two self-adhesive, pre-gelled bipolar surface electrodes (Blue Sensor N, Ambu®, Denmark) were positioned on the muscle body and its distal insertion. A reference electrode was placed on the styloid process of the ulna. EMG signals were sampled at 2 kHz with an online digital notch filter (50Hz), bandpass filtered at 20–450Hz, and amplified using a D360 8-Channel Patient Amplifier (Digitimer®, England).

#### 2.2.4. Accelerometer Recordings

A 8791A250 K-Shear® Miniature Triaxial Accelerometer (Kistler, USA) was secured to the nail of the index finger using micropore adhesive tape (Fig. 1). The accelerometer captured movement in the x, y, and z axes, enabling a complete reconstruction of the motion. The data were collected using a 4-Channel PiezoSmart® (TEDS) Power Supply/Signal Conditioner (Kistler, USA) and recorded as the finger’s acceleration (m/s²). Prior to analysis, the data were low-pass filtered offline using a 20 Hz 4th order Butterworth filter.

**Figure 1:**
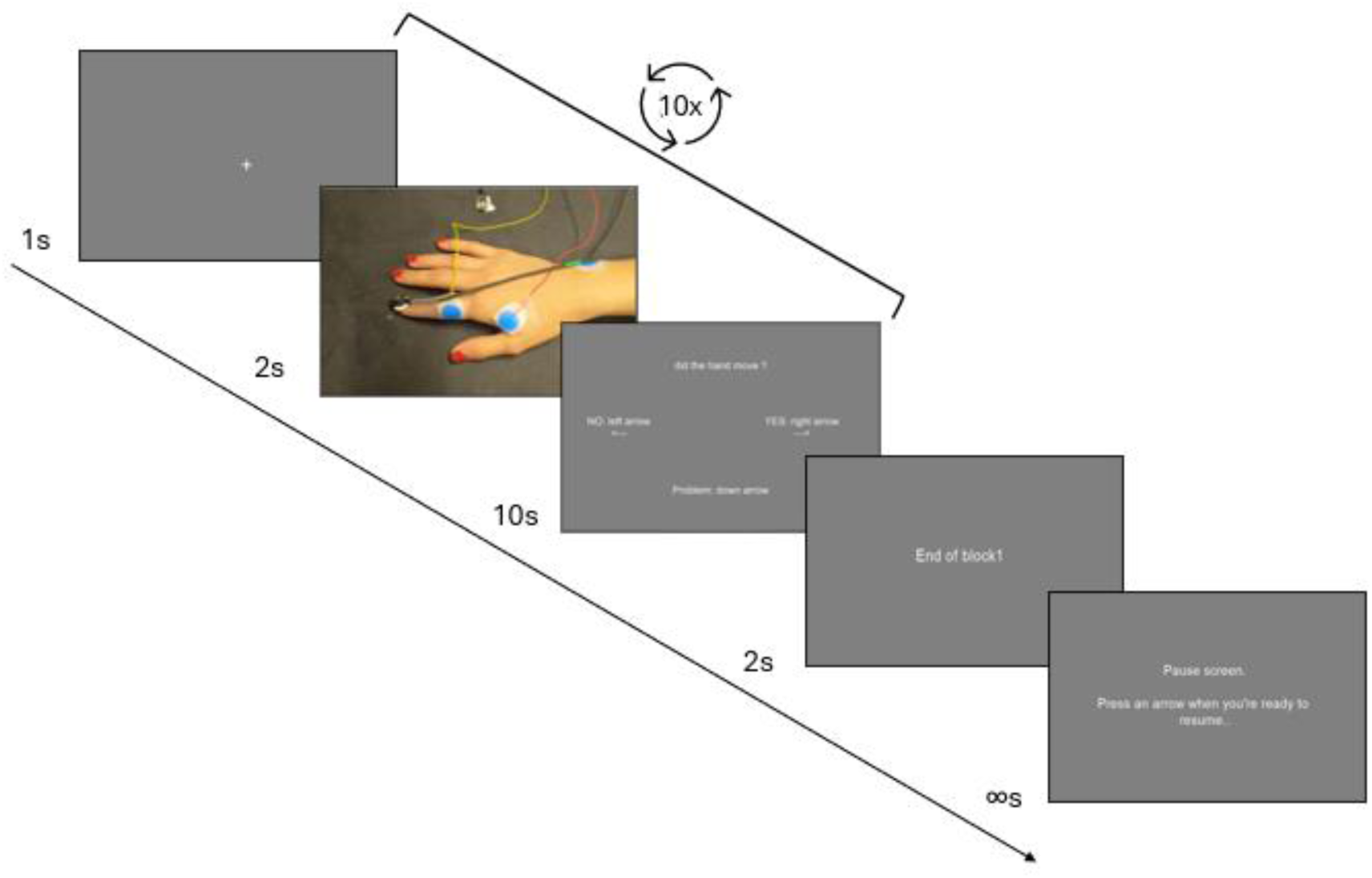
Trial structure for the visual inspection task.

#### 2.2.5. Video recordings

Video recordings of the experiment were captured using a GoPro Hero 8 Black, set to record at 1080p resolution and 120 frames per second. The camera was mounted on a tripod, positioned to film the hand from above and from the contralateral side (see Fig. 1). This setup was chosen to simulate the perspective of an experimenter during a typical experimental session and offered a clear view of the hand.

#### 2.2.6. Protocol Design

Participants sat on a chair with the palmar side of the dominant hand resting face down on the table, and were able to make adjustments until they found a comfortable position that they could maintain throughout testing. The corticomotor area over the left hemisphere corresponding to the right first dorsal interosseous (FDI) muscle was identified, and the motor hotspot was determined using a figure-of-eight coil (2 × Ø70 mm) held tangentially to the scalp and positioned at approximately 45° to the sagittal midline, with the handle pointing posteriorly and laterally to induce a posterior– anterior current in the cortex, which is the standard orientation for FDI hotspotting (51,52). The coil placement is determined by record stimulation sites based on the standard MRI template embedded in the software. A theoretical FDI hotspot was first located within the hand area of the motor cortex. The coil was then systematically moved in 0.5 cm steps in posterior–anterior and medial–lateral directions until the optimal scalp position eliciting the largest and most MEPs were identified. The final hotspot coordinates were stored within the neuronavigation system to ensure consistent coil placement throughout the session. RMT was assessed using EMG, defined as the lowest intensity of maximal stimulator output (MSO) to produce a MEP of ≥50µV in at least 5 out of 10 trials. Stimulation was randomly applied at intensities ranging from −5%MSO to +20%MSO of the EMG-defined RMT. An external experimenter (A.L.) visually monitored the participant’s hand and signaled to the experimenter delivering TMS (G.H.) if an intensity of TMS that produced a movement on each trial (i.e. 10/10 trials) was observed, and future intensities of stimulation were applied only below this intensity. This procedure identified the lowest stimulator intensity above the EMG-defined RMT that consistently elicited visible movements in all trials (i.e., 10/10)(range: +9% to +19% MSO).

#### 2.2.7. Data Processing

EMG and accelerometer data were recorded using a Power1401–3A system and Signal software (both from Cambridge Electronic Design, England). The data were processed using custom MATLAB scripts.

EMG signal analysis focused on identifying the ‘peak-to-peak’ amplitude of the Motor Evoked Potential (MEP), which was calculated as the difference between the highest positive and negative peaks of the response in a post-stimulation window of 200ms.

Accelerometer data, capturing movements in the x, y, and z axes, were combined using the Cartesian formula (absolute acceleration = √ (x² + y² + z²)) to generate a single value representing the total acceleration of the sensor (i.e. a measure sensitive to any movement of the finger in response to TMS, analogous to visual inspection). The derivative of the acceleration was then calculated to determine the ‘jerk’ (m/s³), with the absolute peak jerk being the primary signal of interest. Jerk is particularly useful for analysing rapid or transient movements, which are common in TMS-induced responses.

The processed accelerometer data were also used to identify the RMT as determined by accelerometry. The RMT was defined as the intensity of the maximal stimulator output (MSO) at which at least 5 out of 10 trials exhibited a peak jerk (m/s³) exceeding the 95% confidence interval calculated from the 200 ms preceding the TMS stimulation (49)(i.e., a baseline, motionless period for each participant).

### 2.3. Group 2: Video observations to determine the visual RMT

#### 2.3.1. Participants

Initially, 100 participants were recruited through a combination of in-person sessions (20 participants) and the Prolific online platform (80 participants). However, after data verification and attention checks were considered (see below), a total of 64 participants were retained for analysis. These 64 participants (33 females, 31 males) had an average age of 34.1 years (range: 18–62 years, SD = ±9.4 - for full descriptive data for the final sample see Supplementary Table 1). The study was approved by the Ethics Committee Saint-Luc Hospital, UCLouvain (2020/30JUL/389 - DNMT-TMS – amendment 3), and participants gave informed consent though a computerized form. Participants received compensation of up to 8€ for their time.

#### 2.3.2. Procedure and stimuli

The experimental protocol was implemented using PsychoPy version 2024.4, running on Python 3.10 (53) and transformed to PsychoJS experiment to be hosted online on Pavlovia servers (54).

The experiment began with initial screens that provided instructions to the observer (see supplementary materials - Video instruction). Each trial of the experiment then began with a preparation screen (see Figure 1), displaying a fixation cross in the center of the screen for 1 second. The main stimuli in this experiment consisted of short video clips, each lasting 2 seconds, with a single pulse of TMS being delivered at a specific intensity 1 second into the video. Following the video, participants were given a 10-second response window in which they answered “yes” (right arrow) “no” (left arrow) to indicate whether they observed any movement in the hand. In the case of a technical issue with the stimulus presentation (video freezing, images or sound missing), participants could select the option “problem” (down arrow), which would mark the trial for removal from any subsequent analysis.

After the initial instructions were provided, participants observed two example videos: one without any movement in response to TMS, and one with a pronounced movement in response to TMS. Participants then completed blocks in which they observed 10 trials where TMS was delivered at a fixed intensity of MSO. The intensity assigned to each block was determined randomly by the program, so that the order of presentation did not follow a progressive or predictable sequence. At the end of each block (i.e., after viewing all 10 videos for a given intensity level), participants encountered a pause screen and were able to rest as long as they required before beginning the next block of trials.

Each participant in Group 2 was randomly assigned to view videos taken from one of the participants in Group 1 (from this point onwards we refer to participants in Group 2 as “observers” to help distinguish them from participants in Group 1). Notably, the number of trials required to produce a clear visual response in all trials at a fixed intensity varied from participant to participant in Group 1. As a result, the total number of videos available for observation also differed across participants in Group 2, meaning that not all observers viewed the exact same video set. Importantly, however, all observers completed the full set of videos assigned to them. Observers saw trials delivered across at least 15 intensities, and a maximum 20 intensities (average 17.8 intensities – see also table 1 for summary information). Every 5 blocks, an attention task appeared and asked the observer to press the “up” arrow during both offline or online session; this attention check was designed to be able to catch any participants who were not following the instructions given (e.g. simply repeatedly pressing the same response button across all trials, or randomly selecting one of the three buttons that were normally used to provide a response).

**Table 1:**
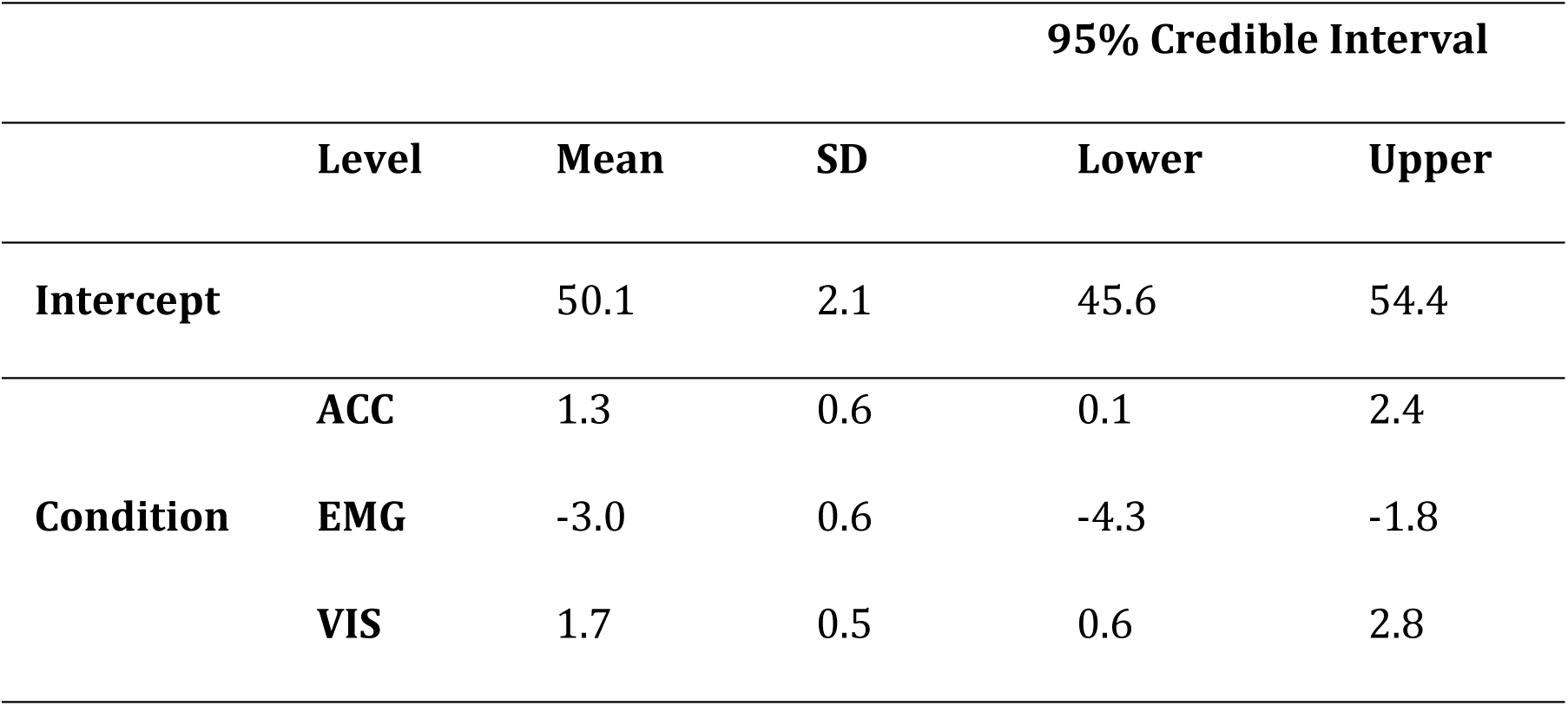
Model-averaged posterior summary statistics for the effects of condition on Resting Motor Threshold (RMT). The intercept reflects the grand mean baseline RMT (in % of maximum stimulator output), while condition estimates indicate the deviation from the intercept for each technique: accelerometry (ACC), electromyography (EMG), and visual inspection (VIS). Positive values represent an overestimation relative to the baseline, and negative values represent an underestimation. Posterior means, standard deviations (SD), and 95% credible intervals are reported.

#### 2.3.3. Data processing

For each observer, the number of “movement” and “non-movement” responses were recorded. These responses were then collapsed across each intensity level to determine the RMT, defined as the intensity at which the proportion of trials with a “movement” response reached or exceeded 50%. This approach is equivalent to the standard EMG/visual method but accounts for any trials that were removed due to technical errors. In total 34 trials were removed (i.e. 0.3% of all trials presented).

### 2.4. Statistical analyses

#### 2.4.1. Bayesian ANOVA

A Bayesian ANOVA was implemented to analyze the effect of the conditions on the dependent variable RMT. This approach allows for the estimation for the effects of conditions (EMG, ACC, and VIS) while accounting for model uncertainty. Random effects, including video series (i.e. different participants from Group 1) and observers (i.e. different participants from Group 2), were incorporated to control inter-individual variability. To interpret the Bayes factors (BFs), statistical analysis followed the classification proposed by Lee & Wagenmakers (55). A grouped scatter plot was created to visualize the data distribution and inter-individual variability in RMT measurements. To further refine the analysis, a Model-Averaged Posterior Summary was conducted, providing weighted mean estimates of condition effects while integrating model selection uncertainty. The posterior mean, standard deviation, and 95% credible intervals were extracted for each condition (EMG, ACC, VIS) to offer a more robust evaluation of their impact on RMT.

Bland-Altman Plots Separate Bland-Altman Plots were generated to examine the agreement between EMG (i.e. the gold standard) and ACC, as well as EMG and VIS, by analyzing systematic biases and limits of agreement. The Bland-Altman plot takes the difference between two measures plotted against their mean, and is therefore useful to observe systematic bias from one method to the other. This visualization was essential for identifying potential measurement discrepancies between the different methods, and assessing their level of agreement. For data examining VIS conditions, we calculated 95% confidence intervals based on a bootstrapping procedure. For each of the video series presented (i.e. data from a single participant in Group 1), 10,000 resamples of the corresponding VIS data (i.e. observers from Group 2 who saw videos from that specific participant in Group 1) were generated with replacement. The mean for each resample was then taken, allowing us to identify a 95% confidence interval (i.e. the 2.5th and 97.5^th^ percentiles). This approach thus ensured that the analyses were not biased by unequal observer contributions or systematic rating sequences, allowing us to obtain more robust estimations of the effects across conditions.

#### 2.4.2. Intraclass Correlation Coefficient (ICC) analyses

Several reliability and agreement analyses were performed to model condition effects. First, Intraclass Correlation Coefficient (ICC) with a two-way random effect model of absolute agreement and single measures (i.e. ICC(2,1), used to evaluate the agreement across raters/measurements) was computed to assess the absolute agreement between EMG (gold standard) and VIS, considering differences across observers(56). Given that the number of observers varied between participants and video series, we implemented a bootstrap-based resampling procedure. We resampled the data 10,000 times, such that each of the 5 different video series (depicting data from Group 1) was assessed by one randomly selected observer (i.e. a single participant from Group 2 was selected from all those available who had watched the corresponding video series). The ICC(2,1) was then calculated for each resample. This approach allowed us to calculate both the mean value, and a bootstrapped 95% confidence interval (taking the 2.5% and the 97.5% percentile ICC values), allowing us to generate a more detailed estimation of the distribution of ICC values. To verify that this resampling did not bias the results, an ICC analysis based on the mean ratings per condition (i.e. ICC (2,k)) analysis with a two-way random effect model of absolute agreement and multiple measures was also calculated and compared with the ICC (2,1) confidence interval.

Additionally, an ICC (2,1) was computed to assess agreement between ACC and VIS conditions, following the same bootstrapping resampling to ensure robustness despite the variability in observer numbers. The ICC (2,k) was computed following the same methodology as described above.

Finally, the ICC (2,1) was computed to assess agreement between ACC and EMG conditions following without bootstrapping due to the number of data acquired (no ICC (2,k) analysis was performed as only one measure was available per participant in Group 1).

To establish reliability thresholds, analysis referred to the classification proposed in Koo & Li (2016) (56) as values below 0.5 indicate poor reliability, those between 0.5 and 0.75 reflect moderate reliability, values ranging from 0.75 to 0.9 suggest good reliability, and values exceeding 0.90 denote excellent reliability.

## 3. Results

### 3.1. Bayesian Model Comparisons

Bayesian Model Comparison of a null model (which included video series and observers as random effects), and a ‘condition’ model (which included the conditions EMG, ACC, and VIS as fixed effects) indicated there was “extreme” evidence in favour of the model including the factor of condition (BF_10_= 11381). A grouped scatter plot demonstrates the effect of the different conditions (Fig. 2).

**Figure 2.**
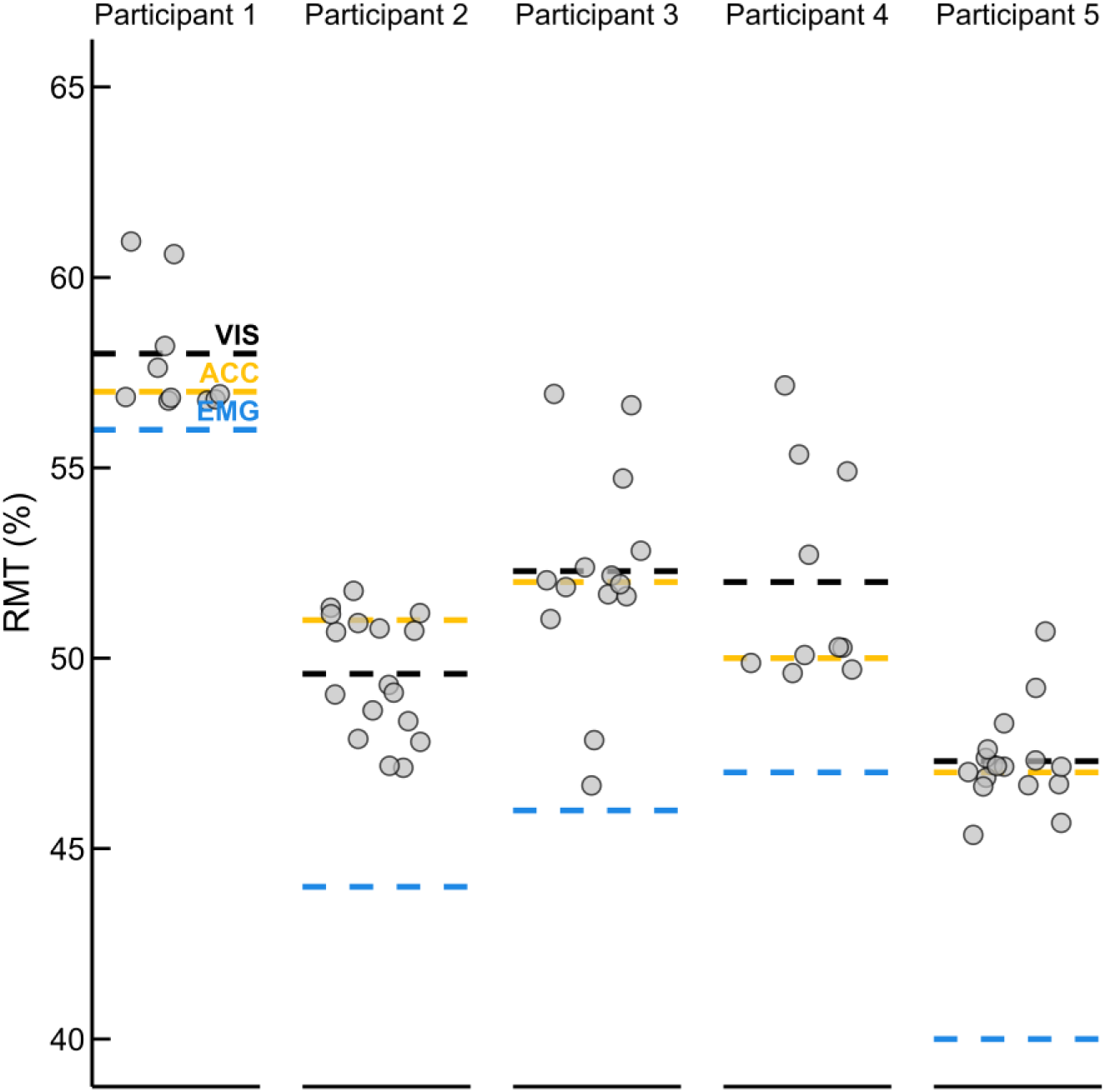
Individual Resting Motor Threshold (RMT) estimates by method and participant. Each panel shows data for one of the five participants from Group 1. Grey dots represent RMT estimates based on visual inspection responses from one of the individual observers in Group 2. Dashed lines indicate the mean RMT obtained via each method: electromyography (EMG, blue), accelerometry (ACC, yellow), and visual inspection (VIS, black).

The Model Averaged Posterior Summary provides estimates for key variables (Table. 1). Notably, the RMT as estimated by EMG was lower than that for either ACC or VIS, with non-overlapping confidence intervals for these comparisons indicating there was a statistically meaningful difference when estimating the RMT using EMG compared to other techniques. By contrast, the overlapping 95% confidence intervals for ACC and VIS indicates their estimates are statistically indistinguishable within the model, suggesting no meaningful difference in estimates of the RMT exists between these two techniques. Furthermore, the positive estimates for ACC and VIS and the fact that the 95% credible intervals remain entirely above zero, suggesting a tendency to overestimate RMT when compared to EMG.

### 3.2. Bland-Altman analysis

When comparing EMG to VIS (Fig 3A), the analysis revealed a systematic bias, as the 95% confidence interval of the mean difference did not include zero, indicating that VIS consistently overestimated RMT relative to EMG rather than fluctuating randomly around it (mean=5.30, 95%CI=2.30-7.22) Additionally, the wide confidence interval suggested substantial variability in the differences between VIS and EMG data.

**Figure 3:**
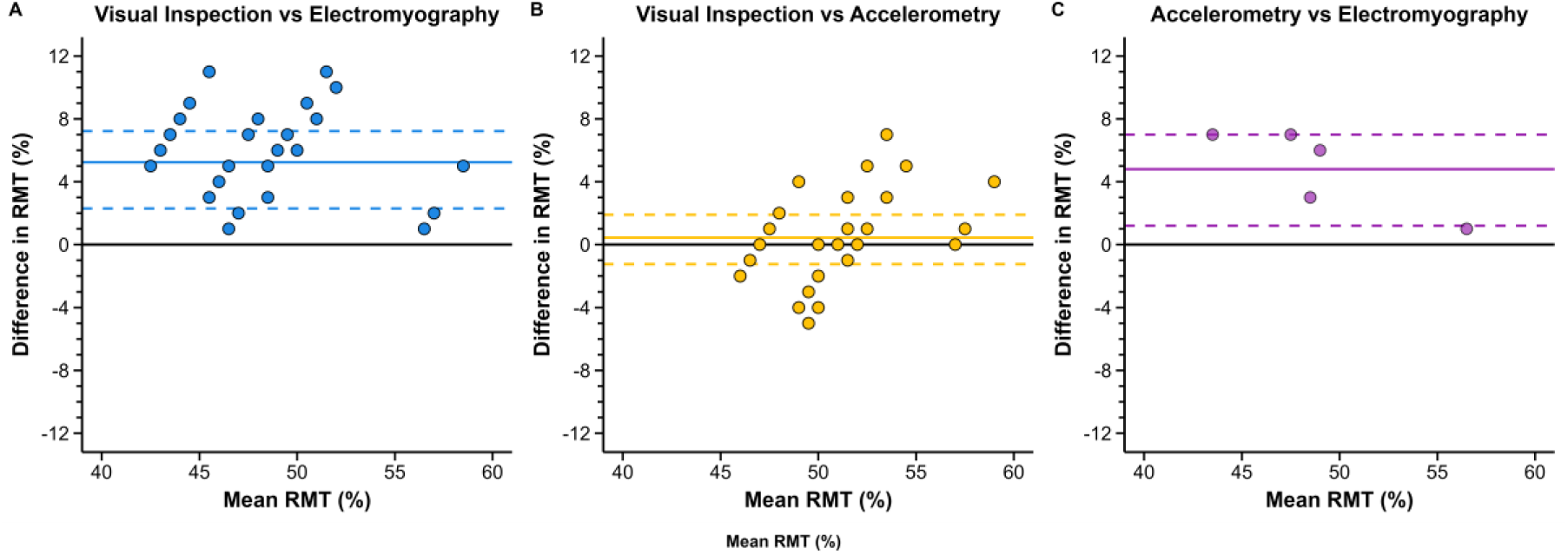
Bland-Altman plots comparing Visual Inspection with Electromyography (A), Visual Inspection with Accelerometry (B) and Accelerometry with Electromyography (C) as reference methods for resting motor threshold. The x-axis represents the mean RMT (%) between the two methods, while the y-axis shows the difference in RMT (%) between visual inspection and the reference method. Each dot represents an individual measurement pair (though note that the data presented here are representative of a bootstrap analysis with 10,000 resamples; due to the high density of data, numerous data points overlap). The solid horizontal line indicates the mean bias, representing the systematic difference between the methods, while the dashed lines denote the 95% confidence interval. Each plot includes 10,000 data points from the resamples. Due to the high density of data, numerous points overlap in regions, leading to visual clustering.

By contrast, when comparing ACC to VIS (Fig 3B), the analysis indicated no systematic bias, as the 95% confidence interval of the mean difference included zero (mean=0.44, 95%CI=-1.25-1.90). However, despite this absence of bias, the broad confidence interval suggested high variability across observers, indicating that while VIS did not consistently over- or underestimate ACC, individual assessments varied considerably between observers.

Finally, comparing EMG to ACC (Fig 3C), the analyses showed systematic bias, as the 95% confidence interval of the mean difference is higher than zero (mean=4.8, 95%CI =1.20-7.00).

### 3.3. Intraclass Correlation Coefficient Analyses

Summary statistics for the bootstrapped analysis are provided in Table 2.

**Table 2:** Intraclass Correlation Coefficient (ICC) values comparing Resting Motor Threshold (RMT) point estimates across methods. ICC (2,1) values reflect agreement between individual observer responses, while ICC (2,k) values reflect agreement based on average ratings per condition. Comparisons include visual inspection (VIS), accelerometry (ACC), and electromyography (EMG). Values are presented with their corresponding 95% confidence intervals. No ICC (2,k) was calculated for the ACC–EMG comparison as only one comparison was available per due to limited sample size.

|  | ICC (2,1) |  |  | ICC (2,k) |  |  |
| --- | --- | --- | --- | --- | --- | --- |
|  | Mean Point | 95% CI |  | Point | 95% CI |  |
|  | Estimate | Lower | Upper | Estimate | Lower | Upper |
| **VIS vs ACC** | 0.845 | 0.615 | 1.000 | 0.980 | 0.815 | 0.997 |
| **VIS vs EMG** | 0.580 | 0.389 | 0.748 | 0.753 | 0.132 | 0.973 |
| **ACC vs EMG** | 0.587 | -0.108 | 0.943 | N/A | N/A | N/A |

For the bootstrap based ICC (2,1) analyses, when comparing VIS to EMG, the mean ICC value identified only ‘acceptable’ reliability, and the 95% confidence interval included responses ranging from ‘unacceptable-to-good’ (mean ICC=0.58, 95%CI=0.39-0.74), with the majority of resamples indicating only ‘acceptable’ reliability. By contrast, when comparing VIS to ACC, the mean ICC value identified ‘good’ reliability, with a 95% confidence interval including a range of responses spanning from ‘acceptable-to-excellent’ (mean ICC = 0.84, 95%CI=0.62-1.00) (Fig. 4).

**Figure 4:**
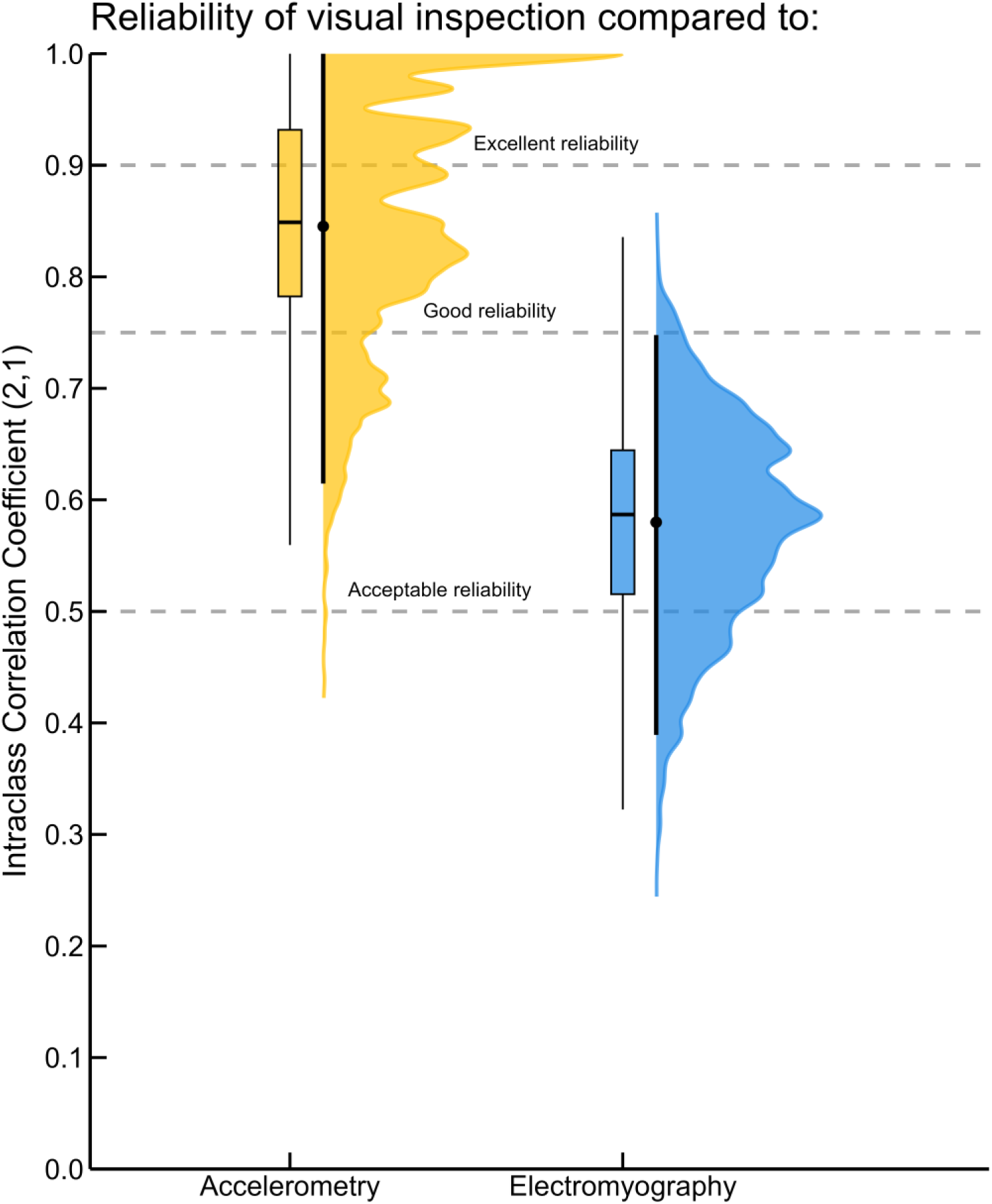
Results of the bootstrap analysis demonstrating the distribution of Intraclass Correlation Coefficients (ICC 2,1) for visual inspection compared to accelerometry (Yellow) and electromyography (Blue). Violin plots represent the full distribution of ICC values with the dot representing the mean and vertical line the 95%CI, while box plots indicate the median, interquartile range, and overall dispersion. Each dashed line represents the reliability threshold based on Koo and Li (2016).

To complement these results, ICC (2,k) values based on averaged ratings were also calculated. These showed “excellent” reliability for VIS vs ACC (point estimate = 0.980, 95% CI = 0.815–0.997) and “good” reliability for VIS vs EMG, though the 95% confidence interval indicated considerable variability, including responses ranging from ‘unacceptable-to-excellent’ (point estimate = 0.753, 95% CI = 0.132–0.973).

Finally, the ICC (2,1) comparison between EMG and ACC showed “acceptable” reliability, again with a wide 95% confidence interval, which included responses ranging from ‘unacceptable-to-excellent’ (point estimate= 0.587, 95%CI=0.108-0.943). No ICC (2,k) was calculated for this comparison due to the limited number of observations available for EMG and ACC.

## 4. Discussion

The present study compared the resting motor threshold as estimated using electromyography, accelerometry, and visual inspection. A series of analyses indicated that while estimates of the resting motor threshold produced by accelerometry and visual inspection are generally similar, these techniques typically over-estimate the threshold as determined using electromyography. However, further analysis supported the view that accelerometry provides a more objective and quantifiable approach to determining the resting motor threshold than visual inspection, which varied considerably between observers. These findings support the view that electromyography remains the gold-standard for estimating the resting motor threshold, but in cases where the use of electromyography is not feasible, the objective and less variable nature of accelerometry provides a preferable alternative to visual inspection.

The present manuscript supports the view that estimation of the resting motor threshold using electromyography represents the ‘gold standard’. Bayesian Model comparison and Bland-Altman analyses both indicated that visual inspection over-estimated the RMT when compared to the use of EMG. This overestimation is easily explained by the fundamental differences in what each method measures. EMG captures an intrinsic characteristic of the muscle—its electrical activity—providing a direct assessment of neuromuscular activation. In contrast, accelerometry and visual inspection rely on an extrinsic characteristic (i.e., overt movement of the finger). Muscle contraction (detectable by EMG) does not always result in displacement of the finger (as measured using ACC and VIS), and movement requires overcoming factors such as inertia and frictional forces. These differences would explain why ACC and VIS tend to overestimate RMT, as they may fail to detect subtle contractions that do not produce sufficient movement.

These results are partially in line with the previous research that has compared the use of visual inspection to EMG when estimating the resting motor threshold (43,44). However, the protocols and findings of these previous studies remain subject to discussion. Pridmore et al. (1998) (42) reported that motor thresholds determined using EMG compared to visual inspection showed a difference of less than 10% in percent of maximum stimulator output, suggesting a general consistency between these approaches. However, they also found that in most cases, the threshold detected through visual inspection was reached at *lower* intensities than with EMG. This discrepancy raises questions about the comparability and sensitivity of these methods in assessing cortical excitability, particularly regarding the potential influence of methodological differences on threshold determination. Meanwhile, Balslev et al. (2007) (45) employed a visual inspection protocol assessing the effects of eight different coil orientations on the resting motor threshold as estimated using EMG and visual inspection. While their results indicated that visual inspection typically over-estimated the RMT as determined by EMG by approximately 2% of MSO, as the comparison between these two methods was not the main focus of their study, only data from four participants using a primarily repeated-measures design (i.e. 8 different coil orientations were assessed for each participant) was collected. This procedure may therefore have led to an under-estimation of differences between the RMT as estimated using EMG compared to visual inspection. We also note that the results of the present study are generally in line with the findings of our previous research (49), in which we found that the RMT as estimated via EMG and ACC were highly correlated, but that RMTs identified by EMG were generally lower than those identified using accelerometry. An important consideration when interpreting our findings is the pronounced measurement error inherent to VIS, which distinguishes it from techniques such as EMG and accelerometry. When the same motor event is recorded and analyzed using EMG or accelerometry, and the same detection algorithm is applied, the outcome is reproducible, yielding a single, objective value per trial. In contrast, VIS introduces considerable variability, as it relies on subjective human judgment.

Further analyses considered whether accelerometry provided any benefits compared to visual inspection. Here we note that while accelerometry provides a single measurement that can be quantified objectively, visual inspection, by comparison, remains a more subjective approach to identifying the Resting Motor Threshold. Bland-Altmann analysis indicated that while the average RMT provided by visual inspection matched well to that of estimates provided by EMG, there was considerable variability around the mean (95%CI ranging from –4 to +5 MSO). Similarly, our ICC bootstrapping analysis indicated that while on average VIS has “good” reliability when compared to ACC, there was again a large disparity in the 95%CI, indicating that, in the best-case scenario, an observer (VIS) may provide a value close to ACC, but in other cases, their estimate of the RMT could be far from it.

Given these findings, we propose that researchers who are not able to use EMG to determine the RMT should prefer the use of accelerometry, rather than visual inspection, wherever possible. Unlike VIS, which depends on individual observers and introduces variability, accelerometry provides a single, consistent measurement per trial, reducing subjectivity and improving measurement reliability. Nonetheless, a limitation of the accelerometry is that it requires real-time (online) data processing through a dedicated script.

However, several limitations should be acknowledged. First, the sample size for the EMG and the accelerometry conditions was limited to five participants, which constrains the generalizability of these findings and increases the risk of sampling bias. We note, however, that the results of the comparisons between EMG and ACC in the present study are in agreement with our previous work which has demonstrated the same effects in a larger sample size (49). Second, a large proportion of the data for visual inspection were collected in a remote, online experiment, which prevented close monitoring of participant engagement. The inclusion of attention checks therefore helped to maintain high quality data (i.e. approximately 1/3 of participants were rejected based on this measure). Notably, no participants were rejected in the in-person condition, suggesting that the physical presence of the experimenter likely contributed to sustained engagement and compliance with the task. Third, the visual inspection condition relied on a participant reviewing recorded videos, rather than observing the stimulation in real time.

A further limitation is that the observers in the present study were relatively ‘naive’ compared to experienced TMS experimenters. While training might reduce variability, individual perceptual differences would likely persist, meaning that visual inspection would still be less reliable than accelerometry. However, VIS could be considered in studies using ‘main’ protocols delivered at submaximal RMT-based intensities (e.g. 90% of RMT) to avoid safety issues (Rossi, Simone et al., 2009),

In addition, the recorded hands used for Group 1 were limited to a small sample (n=5), which restricted the diversity of visible features such as gender, skin tone, and hand or digit morphology. As a result, observer–actor mismatching may have contributed, even in a small part, to the perceptual judgments of finger twitches. While unlikely to account for the main effects observed, this highlights the need for future studies to examine visual detection of MEP responses across more demographically diverse samples.

Thus, visual inspection would still introduce variability to the measurement of the resting motor threshold that accelerometry would remove. Additionally, the convergence of our findings with previous research (e.g., our prior work on EMG and ACC correspondence) further supports the reliability of the present data. In particular, the strong agreement observed between EMG and ACC measurements confirms the replicability of our previous results, and highlights the potential of using an equation to convert the RMT as calculated by accelerometry to an estimated value of the RMT as would be determined by EMG. Accelerometry therefore remains a viable alternative when EMG is not available.

In conclusion, the present study indicates that EMG provides the most sensitive approach to identifying the RMT, most likely due to its ability to measure the intrinsic excitability of the corticospinal system. In situations where access to electromyography is limited, researchers should consider using accelerometry in the place of visual inspection, as accelerometry provides a more objective and reliable approach.

## 5. Research data

The research data supporting the findings of this study are stored in an OSF (Open Science Framework) repository and are available upon reasonable request from the corresponding author at. Open public access is restricted to protect participant confidentiality in accordance with the guidelines of the approving ethics committee.

## 6. Acknowledgements

Gautier Hamoline is supported by a PhD fellowship from the Faculty of Motricity Sciences, UCLouvain, and the Belgian Fund for Scientific Research (FNRS J.0084.21). Robert Hardwick is supported by grants from the UCLouvain Special Research Funds Seedfunds (FSR 1C21300057) and the Belgian Fund for Scientific Research (FNRS F.4523.23 and FNRS J.0084.21). Marcos Moreno-Verdu is supported by a grant from the Belgian Fund for Scientific Research (FNRS 1.B359.25). Elise Van Caenegem is supported by a grant from the Belgian Fund for Scientific Research (FNRS 1.AB19.24). Baptiste Waltzing and Siobhan McAteer are supported by the Belgian Fund for Scientific Research (FNRS F.4523.23).

## 7. Author Roles (CRediT)

| Author | Conceptualization | Data curation | Formal analysis | Funding acquisition | Investigation | Methodology | Project administration | Resources | Software | Supervision | Validation | Visualization | Writing – original draft | Writing - review and editing |
| --- | --- | --- | --- | --- | --- | --- | --- | --- | --- | --- | --- | --- | --- | --- |
| Hamoline<br>Gautier | x | x | x |  | x | x | x | x | x | x | x | x | x | x |
| Antoine<br>Lunardi |  | x |  |  | x |  |  |  |  |  |  |  | x |  |
| Marcos<br>Moreno-Verdu |  |  | x |  |  |  |  |  | x |  |  | x |  | x |
| Baptiste M<br>Waltzing |  | x |  |  |  |  |  |  |  |  |  |  |  | x |
| Elise E Van<br>Caenegem |  | x |  |  |  |  |  |  |  |  |  |  |  | x |
| Siobhan<br>MacAteer |  |  |  |  |  |  |  |  | x |  |  |  |  | x |
| Robert M<br>Hardwick | x |  |  | x |  | x | x |  | x | x | x |  | x | x |

## Supplementary materials

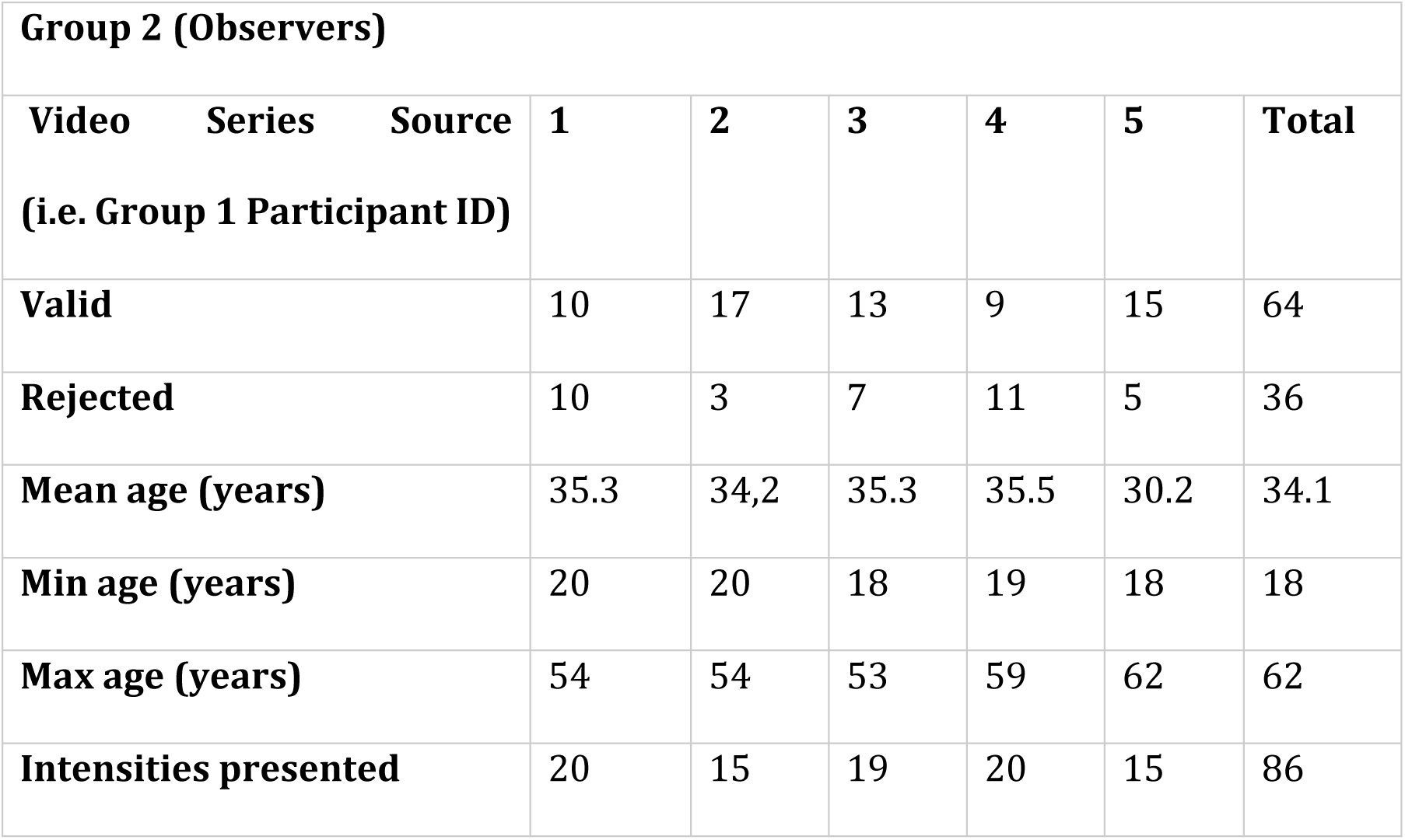

### Descriptive Statistics

|  | RMT |  |  |  |  |
| --- | --- | --- | --- | --- | --- |
|  | 1 | 2 | 3 | 4 | 5 |
| Valid | 11 | 17 | 14 | 10 | 17 |
| Missing | 0 | 0 | 0 | 0 | 0 |
| Mean | 57.9 | 49.6 | 52.3 | 52.0 | 47.3 |
| Std.<br>Deviation | 1.6 | 1.6 | 2.8 | 2.7 | 1.3 |
| Coefficient<br>of<br>variation | 0.027 | 0.033 | 0.053 | 0.053 | 0.027 |
| Range | 4 | 5 | 10 | 7 | 6 |
| Minimum | 57 | 47 | 47 | 50 | 45 |
| Maximum | 61 | 52 | 57 | 57 | 51 |

### Video instructions

**Screen 1**

Hello,

Thank you for participating in this experiment.

The purpose of this experiment is to evaluate the ability of humans to discriminate hand movement during transcranial magnetic stimulation.

your data will be identified with your Prolific ID or a random number generated if you don’t fill it

If you understand, press one of the arrow keys. If you have any questions, contact and press ESC.

**Screen 2**

Each video takes place as follows:

- A cross will appear for one second to allow you to be ready

- The video starts and you will hear a clicking sound (turn on the sound) and a flashlight. This click is the stimulation signal, and is part of the experience.

- After the video, a window will appear, asking you to answer whether your hand has moved or not.

No = left arrow <--

Yes = right arrow -->

You have 10 seconds to respond

If a video has a minor problem respond if your able too,

if not = down arrow

There are 20 blocks of 10 stimulations, after each block, there will be a pause screen to allow you to take a breath.

! There doesn’t have to be parity in the answers. the videos presented to you are randomly chosen.

If you understand, press one of the arrow keys. If you have any questions, contact and press ESC.

**Screen 3**

You will be given two examples to prepare you for the experience.

**Screen 4**

The first video was an example of a moving hand, the second one was a non-moving hand.

Let’s get started.

If you understand, press one of the arrows. If you have any questions, contact and press ESC.

